# A ChIC solution for ChIP-seq quality assessment

**DOI:** 10.1101/2020.05.19.103887

**Authors:** Carmen Maria Livi, Ilario Tagliaferri, Koustav Pal, Endre Sebestyén, Federica Lucini, Andrea Bianchi, Sara Valsoni, Chiara Lanzuolo, Francesco Ferrari

## Abstract

Despite the widespread adoption of the ChIP-seq technique, there is still no consensus on quality assessment procedures. Quantitative metrics previously proposed in literature are not always effective in discriminating the success or failure of an experiment, thus hampering objectivity and reproducibility of quality control. Here we introduce ChIC, a new framework for ChIP-seq data quality assessment that overcomes the limitations of previous solutions. ChIC is the first method for ChIP-seq quality control directly considering the enrichment profile shape, thus achieving good performances on ChIP targets yielding sharp and broad peaks alike. We integrate a comprehensive set of quality control metrics into one single score reliably summarizing the sample quality. The ChIC score is based on a machine learning classifier trained on a compendium with thousands of ChIP-seq profiles, which can also be used as a reference for easier evaluation of new datasets. ChIC is implemented as a user-friendly R/Bioconductor package.

## INTRODUCTION

Chromatin immunoprecipitation followed by high-throughput sequencing (ChIP-seq) is a widely adopted technique for genome-wide mapping of transcription factor (TF) binding sites and chromatin marks distribution [1]. Variability in data quality can be introduced by several factors, such as antibody quality [2] or various experimental protocol steps [3, 4], and may cause substantial differences in the enrichment of the signal compared to the background [5–7]. Quality control (QC) is therefore an important analysis step. Several metrics and guidelines have been proposed [3, 4, 8] but there is still no consensus on QC procedures. Among the many tools for ChIP-seq analysis that have been introduced over the years [6, 9, 10], including peak callers [11–15] or complete analysis pipelines [16, 17], only a few of them report quality metrics. The QC is often limited to generic read-level statistics on the FASTQ file [18] or to selected metrics [10, 19, 20] that do not provide a completely generalizable solution.

Indeed, a critical challenge in quality assessment is the variety of ChIP targets yielding enrichment profiles with distinct characteristics. Namely, different chromatin marks have peculiar distributions along the genome, most notably resulting in peaks with sharp or broad shape in the ChIP-seq enrichment profile [1]. For this reason, they may not be compliant with general guidelines and often have to be reconsidered on a case-by-case basis, as explicitly discussed also in the reference article on this topic by the ENCODE consortium [3]. Consequently, quality assessment is still a partially subjective operation, influenced by the experience of the data curator.

Moreover, a common view shared by literature in the field [4, 7, 21] is that a single score providing a reliable summary of the quality of a ChIP-seq sample would be convenient for end users. Despite a few solutions have been proposed [4, 7, 21, 22], there is not yet a consensus in literature on an unbiased single score to reliably discriminate between good and poor-quality samples. Thus, a broad ensemble of parameters must be considered [3].

To address these limitations, we propose a new **<u>ChI</u>**P-seq quality **<u>C</u>**ontrol framework named “ChIC”. The key rationale of our framework is that the expected shape of the enrichment profile must be considered while assessing ChIP-seq data quality. For this reason, we introduce a new set of metrics, the Local enrichment profile Metrics (LM), by defining quantitative scores that describe the shape of ChIP-seq enrichment profiles (see methods). These metrics are not replacing, but instead extending previously proposed quantitative metrics for scoring ChIP-seq data, that are also computed as part of a comprehensive set of quantitative QC-metrics including ENCODE proposed metrics (EM) and metrics describing the global enrichment (GM) in the ChIP-seq experiment. This comprehensive set of metrics is leveraged to build a machine learning classifier to obtain a single score reliably summarizing the data quality and discriminating between good and bad quality ChIP-seq samples.

To the best of our knowledge, this is the first ChIP-seq QC method directly considering the shape of enrichment, the first one translating the shape of enrichment into quantitative metrics and the first based on machine learning. We also extensively benchmark our method against previously proposed solutions achieving consistently high performances. Finally, ChIC is implemented as a user-friendly R package, compliant with Bioconductor strict standards of stability and documentation. The latest stable version is available at https://bioconductor.org/packages/devel/bioc/html/ChIC.html.

## RESULTS

### Standard QC-metrics are biased by the shape of ChIP-seq enrichment profile

We analysed a large set of ChIP-seq experiments with paired input control, for a total of 3936 samples from large public databases (Table 1), including the ENCODE project [23] and Roadmap Epigenomics Consortium (Roadmap) [24], to build a reference compendium of QC-metrics, including previously proposed and novel metrics (Fig. 1a).

**Table 1.** Dataset description.

|  | **Encode**<br>[23, 28] | **Roadmap**<br>[24] | **Simulated failed**<br>**samples** |
| --- | --- | --- | --- |
| ChIP | 1824 | 1546 | 3370 |
| Control/Input | 344 | 222 | - |
| DNA-binding<br>proteins | 158 | 0 | - |
| Chromatin marks | 12+RNAPol2 | 31 | - |

**Figure 1:**
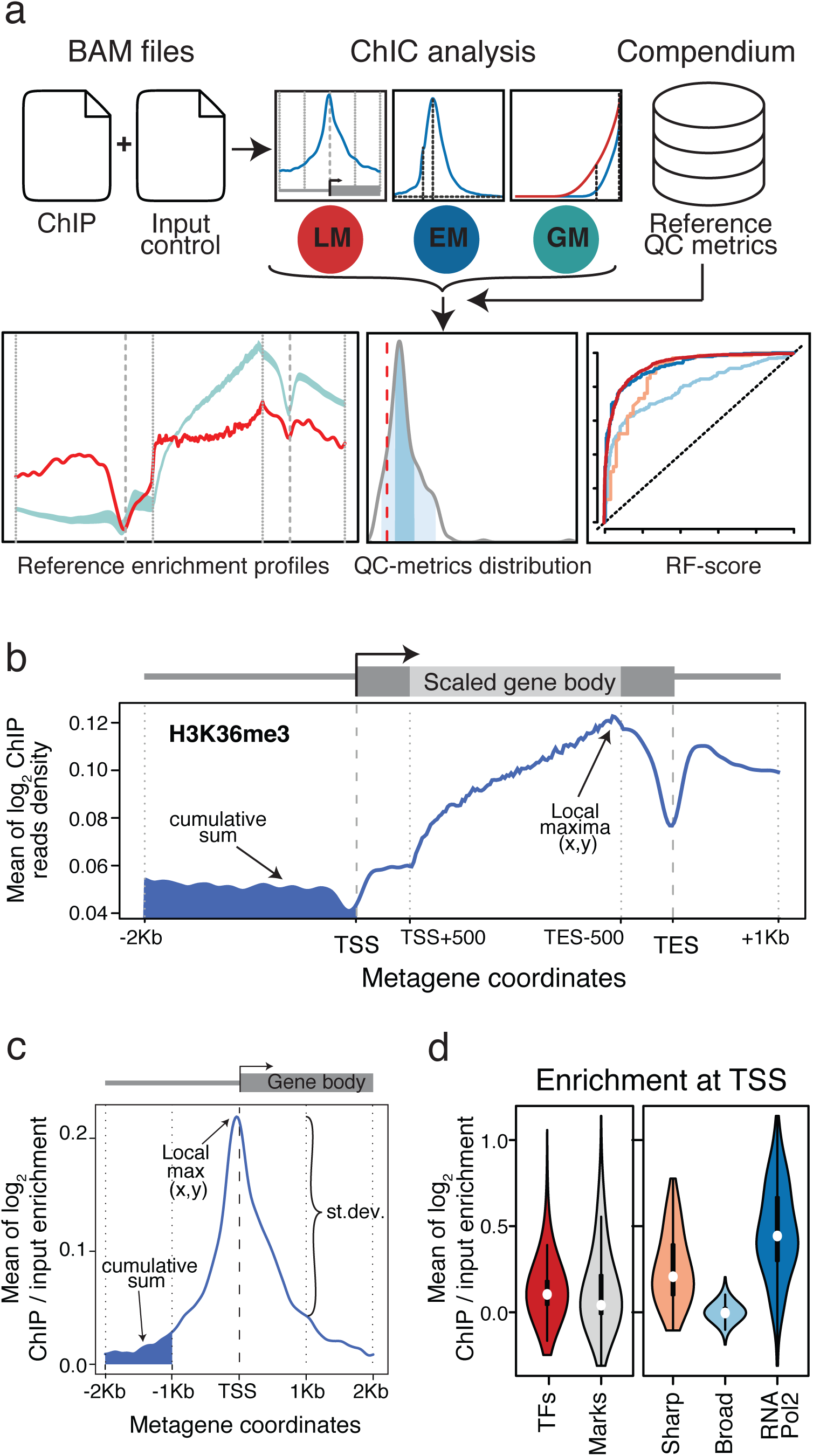
Overview of ChIP-seq quality Control (ChIC) framework and QC metrics. (**a**) Schematic workflow of the ChIP-seq quality Control (ChIC) framework. ChIC implemented as a Bioconductor R package takes as input aligned read files (.bam) for ChIP and control (input) for a query sample. A comprehensive set of QC-metrics is computed, including Local enrichment profile Metrics (LM), ENCODE Metrics (EM) and Global enrichment profile Metrics (GM). The query sample can then be compared to the compendium of reference values in three ways: (i) plotting the metagene profile against the expected profile of the respective chromatin mark class or sub-type; (ii) comparing an individual selected QC-metric against the reference distribution of values of the compendium; or (iii) computing a single score (RF-score) summarizing the sample quality based on our random forest models. (**b**) Example of a scaled whole gene metagene profile for chromatin mark H3K36me3, showing the average ChIP only signal around the TSS, TES and along the entire gene body of annotated genes. The scaled (light grey) and unscaled (dark grey) portions of the metagene profile are marked in the schematic gene model above the plot. The metagene coordinates on the x-axis show the genomic distance in real scale for 2Kb upstream of the TSS and 1Kb downstream from the TES. The signal in the gene body is scaled except for the first and the last 500bp of the gene body. The log_2_ transformed library size normalized ChIP reads density signal is shown on the y-axis. **(c)** Example of TSS centred (unscaled single-point) metagene profile with the normalized (ChIP over input) enrichment signal for all annotated genes for a representative CTCF sample. The TSS and the beginning of the gene body are marked by a dark grey gene model above the plot. The metagene coordinates on the x-axis show the genomic distance (in bp) from the TSS with up- and down-stream regions in real scale, whereas the log_2_ transformed normalized (ChIP over input) enrichment signal is shown on the y-axis. (**d**) LM example: the violin plots show the distribution of the average log_2_ transformed normalized (ChIP over input) enrichment signals taken at the TSS position. ENCODE and Roadmap data samples are grouped by TFs (1125) and chromatin marks (2245) categories in the left plot. The right plot shows instead the values distribution for chromatin marks divided into 3 peak profile classes: Sharp (941), Broad (1177) and RNAPol2 (127, containing different subunits).

Previously proposed QC quantitative metrics for ChIP-seq do not explicitly take into account the shape of the enrichment profile, yet this is affecting the QC score values. For example, some of the QC-metrics proposed by the ENCODE consortium are more effective on ChIP targets yielding narrow peak profiles, such as TFs, because they are designed for ChIP-seq enrichment profiles with sequencing reads localized in small regions. This is the case for all the metrics based on strand-shift analyses [3, 4, 7, 10, 11]. Strand-shift analyses, such as the “cross-correlation” analysis (Supplementary Figure S1a; methods), aim to detect the clustering of reads in a ChIP-seq sample without relying on peak calling. The strand-shift profile is calculated using the position of reads on positive vs negative strand and shifting them towards each other. At each progressive shift the correlation [3, 4] between the position of reads on the two strands is calculated. In other variants of this analysis, other measures like the Jaccard Index [7] or the Hamming distance [15] have been proposed instead to quantify the clustering of reads on positive and negative strands. Then, multiple QC-metrics for the strength of the ChIP enrichment can be derived from the resulting cross-correlation profile, for example the normalized or relative strand coefficient (NSC and RSC, respectively) or the quality control tag (QC tag) [3, 4, 11]. We refer to this set of metrics, along with others described by Landt *et al.* [3], as ENCODE Metrics (EM) (see methods). Notably, the relative strand coefficient (RSC) is proportional to the level of clustering of reads, i.e. the strength of the enrichment signal, in the ChIP-seq samples [3, 4] (Supplementary Figure S1a). As such, it generally assigns higher scores to ChIP-seq profiles with localized enrichment signal (such as TFs), rather than other chromatin marks yielding broader enrichment profiles (Supplementary Figure S1b - left). Indeed, a more detailed grouping of the chromatin marks based on their expected profiles (Sharp, Broad and RNAPol2; Table 2) confirms that chromatin marks with sharp peaks (Sharp), *i.e.* more similar to TFs, get higher RSC scores than marks with broader enrichment profiles (Broad) (Supplementary Figure S1b - right). The mixed profile characteristics of RNA Polymerase 2 (RNAPol2) are reflected as well in the distribution of the RSC metric, showing an intermediate range of values. Thus, relying exclusively on RSC, or its discretized counterpart named QC tag [4], as a single score to select high quality data, may result in discarding good quality samples for chromatin marks with broader enrichment profiles. This kind of bias affects also other frequently used EM scores that directly use sharp peaks to compute the score. This is the case of the fraction of reads inside ChIP-seq peaks (FRiP) [3, 10]. To this concern, we also considered an alternative FRiP_broadPeak metric (Supplementary Figure S1c), to capture features of broader enrichment profiles.

**Table 2.** Grouping of chromatin marks by peak profiles.

| **Binding pattern** | **Chromatin mark** |
| --- | --- |
| Sharp | H3K4me2/3, H3K79me1, H3K27ac, H2BK12ac, H2BK120ac, H2BK15ac, H2BK20ac, H3K4ac, H4K12ac, H4K8ac, H4K5ac, H2AK5ac, H2A.Z/H2AFZ, all remaining acetylations |
| Broad | H3K4me1, H3K79me2, H3K36me3, H3K9me1/2/3, H3K23me2, H3K27me3, H4K20me1, H3K56ac, H3T11ph |
| RNAPol2 | RNAPol2 |

Other methods for assessing ChIP-seq data quality capture features of the global read distribution. An example of this are metrics based on the fingerprint plot, *i.e.* the direct comparison of the cumulative distribution of reads across genomic bins in the ChIP and control samples [8, 22] (Supplementary Figure S1d). Hereafter we are referring to the quantitative metrics that can be derived from the fingerprint plot as Global enrichment profile Metrics (GM) (see methods) [8, 22]. In principle, they are applicable to both TFs and broader ChIP-seq profiles. However, even these metrics examining the global reads distribution do not explicitly account for the expected shape of the enrichment profile expected with different ChIP-seq targets. For example, the “fraction of reads in the top 1% bins” is expected to be higher for TFs as narrow peak profiles accumulate a larger fraction of reads in a limited number of genomic bins. A lower value is expected for chromatin marks that cover larger genomic regions and even lower for the input control samples, where the reads are distributed throughout the genome. However, we did not observe a large difference between values for TFs and other chromatin marks (Supplementary Figure S1e - left plot). This is due to the heterogeneous nature of chromatin mark profiles. Indeed, after grouping them into distinct subclasses (Supplementary Figure S1e - right plot) we observed higher values for Sharp as opposed to Broad, as expected. RNAPol2 is an exception again, due to its mixed peak profile.

Overall, these results confirm the need to account for the different classes of ChIP targets while performing QC assessment. However, the fundamental differences in the peaks shape between distinct profile classes are not directly considered in previously proposed QC metrics. Therefore, it is highly recommended to consider the binding characteristics of the ChIP-seq target when applying metrics for quality assessment.

### Accounting for local enrichment profile shape in QC-metrics

We introduce a novel set of QC-metrics (LM) to achieve a measurable and reproducible assessment of the local enrichment profile shape. More specifically, we use the so-called “metagene” profiles, *i.e.* the average ChIP-seq signal over a set of genes, to derive a total of 214 quantitative features (Fig. 1b, c) (see methods). These include for example the average level of signal enrichment at the annotated transcription start sites (TSS) (Fig. 1d) or annotated transcript end (TES) (see methods). LM values provide important information about the signal distribution and depend not only on the quality of the data but also on the type of chromatin mark under investigation. For example, Histone 3 Lysine 4 tri-methylation (H3K4me3) or TFs usually show marked enrichment at promoter regions, whereas Histone 3 Lysine 36 tri-methylation (H3K36me3) is enriched over the body of transcribed genes (Fig. 1b). LM are intended to capture chromatin mark-specific enrichment features that are instead neglected by previously proposed QC metrics.

### Summarizing ChIP-seq quality in a single QC-score

In order to obtain a single score to reliably summarize the quality of ChIP-seq samples, we built a machine learning classifier. Our compendium of reference quality metrics, including LM, EM and GM, is based on 1125 TF and 2245 chromatin mark ChIP-seq samples, in addition to their input control samples, each described by a total of 245 quantitative features (QC-metrics). Some collected features are expected to show a certain degree of correlation based on how they were defined. To remove uninformative correlating features, the QC-metrics were filtered by using hierarchical clustering based on pairwise distances, then cutting the dendrogram into sub-clusters. Only the most representative element for each cluster (medoid) was retained (see methods; Fig. 2a, Supplementary Figure S2a). The reduced set of features used to train the classifier were defined separately for chromatin marks and TFs. Principal component analysis confirmed that our reduced set of features can capture differences between distinct chromatin mark sub-types (Supplementary Figure S2b).

**Figure 2:**
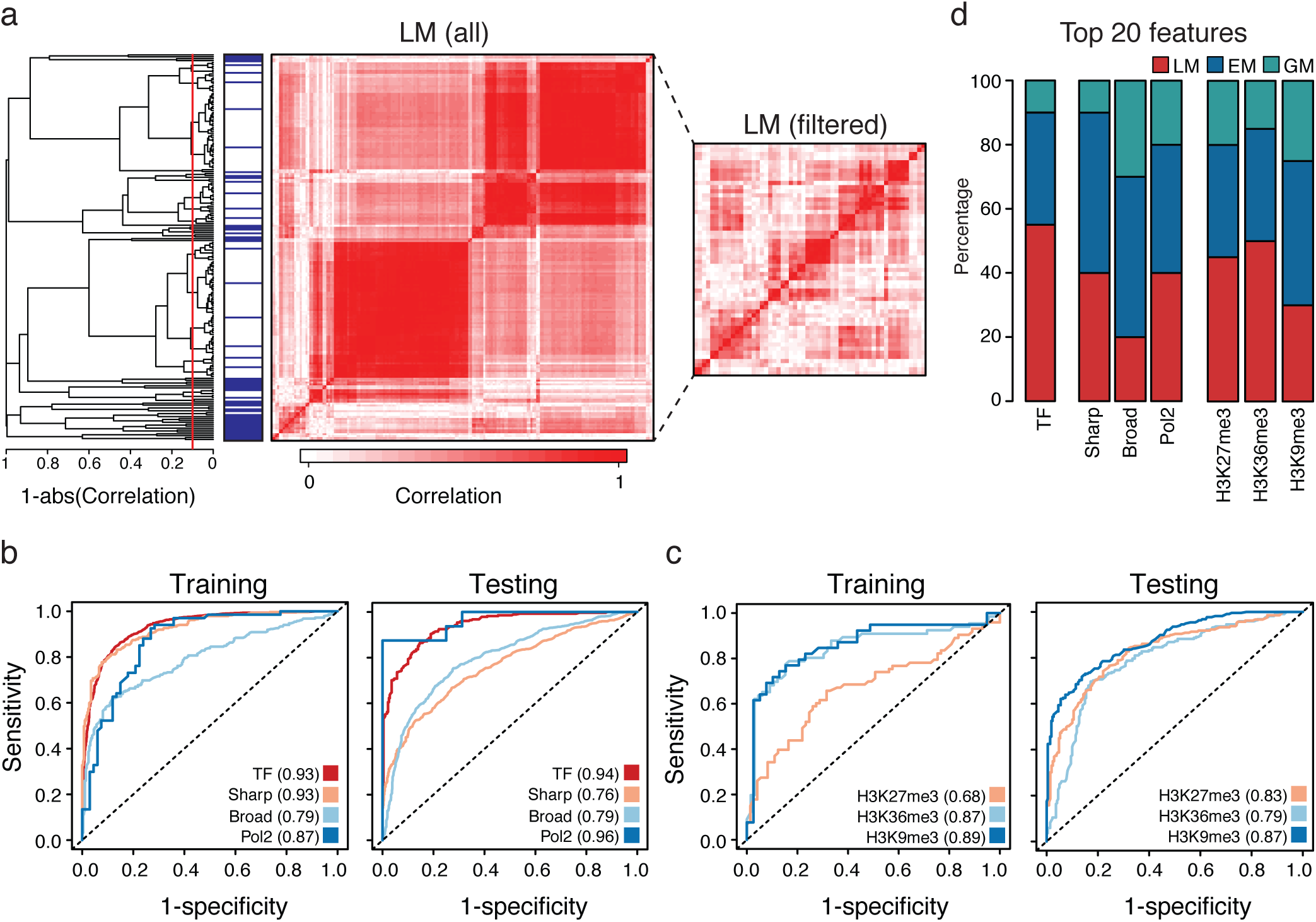
Random forest classifier to assess data quality. (**a**) Correlated features are filtered applying hierarchical clustering on each set of QC-metrics (see methods). The dendrogram shows the result of the complete linkage clustering on LM for chromatin marks with distance measured as 1 - (absolute value Pearson correlation) (EM and GM are shown in Supplementary Figure S2a). The dendrogram is cut at 0.1 and for each resulting cluster the medoid (blue mark in the sidebar) is retained in the set of filtered features. The left heat map shows the pairwise Pearson-correlation between all LM features (before filtering) whereas the right side heatmap summarizes the correlation of the reduced set of features after filtering. (**b**) ROC curves summarizing the performance of each random forest model (TF, Sharp, Broad and RNAPol2) during training and testing with the respective AUCs indicated in the colour legends. The sensitivity is shown on the y-axis and 1-specificity on the x-axis and. The models are trained on ENCODE data (left plot; TF 1125, Sharp 287, Broad 284, RNAPol2 67, only subunit 2A) and independently tested on Roadmap samples (right plot; Sharp: 590, Broad: 892). TFs and RNAPol2 testing data are taken from left-out ENCODE samples as Roadmap does not provide TF and RNAPol2 data. The negative class samples are in both cases their respective simulated problematic counterpart (see methods). (**c**) Similar to panel **b**, ROC curves showing the performance of the random forest models specific for H3K9me3, H3K27me3 and H3K36me3 histone marks, during training (left plot; 39, 73 and 66 samples, respectively) and testing (right plot; 219, 209 and 221 samples, respectively). (**d**) The 20 features with highest importance value in each random forest model (x-axis labels) are grouped by category (LM, GM, EM - colour legend). The relative frequency of each category is reported as percentage in the stacked barplot (y-axis).

The best practice would require a large and numerically balanced set of true positive and true negative quality samples to train the machine learning classifier, as well as to quantitative benchmark ChIP-seq quality control metrics by building ROC curves. Therefore, we synthetically generated the true negative quality dataset by simulating lower signal-to-noise ratio in the ChIP samples. For this purpose, we down-sampled 60% of the reads located within a coarse grain definition of regions with ChIP signal enrichment over input (see methods). It should be remarked that we do not apply uniform subsampling, but we specifically subsample only the ChIP sample reads located within regions of signal enrichment (broad peaks - see methods), while leaving the coverage of the background (noise) regions unchanged. This procedure allows simulating data of lower quality, *i.e.* lower signal-to-noise ratio as expected in a problematic experiment, while at the same time preserving some key features in the distribution of signal in real experimental data. With this approach, we created a problematic counterpart for each sample from the ENCODE and Roadmap datasets, to have an equal number of samples in the positive and negative sets.

Then, random forest models were trained in a 10-fold cross-validation on the ENCODE dataset and independently tested on Roadmap data to exclude circularity with the selected features. In total, we trained 4 separate models: one for TFs and three for chromatin marks (Broad, Sharp and RNAPol2). The positive prediction score obtained from the random forest classifier can be used as a single QC-score assessing the quality of ChIP-seq samples, that we named ChIC RF-score. In the 10-fold cross-validation during training, the models obtained good prediction rates with an area under the curve (AUC) equal to 0.79 for broad marks and 0.93 for sharp marks or TFs, considered separately (Fig. 2b). The independent testing on Roadmap data confirms the predictive stability of the trained models. As RNAPol2 and TFs are not available in the Roadmap dataset, the independent testing was performed on a left-out subset of the ENCODE data for these two profile classes. The optimal thresholds on the RF-score values for discriminating good and poor-quality samples can be defined based on the ROC curves and are reported in (Supplementary Table S1). As a further control we verified that the down-sampling procedure adopted to simulate lower signal-to-noise ratio is not biasing the results, as the RF-score is not correlated to sequencing depth *per se* (Supplementary Figure S2c).

We further reasoned that broad chromatin marks include profiles that can be very different in shape and distribution, as they are associated to different biological mechanisms like Histone 3 Lysine 9 tri-methylation (H3K9me3), associated to heterochromatin; Histone 3 Lysine 27 tri-methylation (H3K27me3), associated to Polycomb Repressive Complex 2 (PRC2) regulation; and H3K36me3, associated to the body of transcribed genes. We tested the performance of scoring samples for these 3 individual marks by training separate random forest models (Fig. 2c, Supplementary Table S1). These more specific random forest models were able to further improve the AUC for H3K9me3 (0.89) and H3K36me3 (0.87) for ChIC RF-score, compared to the AUC for all Broad marks considered together (0.79). For H3K27me3 instead the specific model yielded a 0.68 AUC, thus not improving the classifier performance, although it yielded a higher AUC (0.83) in the testing set. These results show the importance of stratifying chromatin marks by their peak profile class to assess their quality.

To understand the contribution of individual QC-metrics to the ChIC RF-score, we analysed the feature importance provided by each random forest model (Supplementary Figure S2d). Among the 20 most important features we find several LM values, especially for individual broad marks (H3K27me3, H3K9me3), for RNAPol2 and TFs (Fig. 2d) and some widely used EM and GM, such as the FRiP or the fraction of reads in the top 1% bins. This is in line with the fact that these features are indeed valuable and established parameters to assess ChIP-seq sample quality that are effectively complemented by LM scores. Notably, some features related to input control sample rank among the top ten most important ones, like the “fraction of reads in top 1% of bins” for the input control sample in the sharp peaks features ranking (Supplementary Figure S2d). This is confirming the importance of the characteristic of the paired input control data, which is implicitly taken into account as well in our comprehensive set of metrics [25]. Globally, these results indicate that it is crucial to incorporate both local and global features to assess the quality of the sample.

### Benchmarking against other ChIP-seq QC scores

To assess more thoroughly the performance of ChIC RF-score, we benchmarked it against other individual quantitative scores to assess ChIP-seq QC, including independent metrics not comprised in our compendium. Namely, in the benchmarking we considered: 1) the RSC and NSC values, derived from cross-correlation strand-shift analysis as described in [3], which are also part of the EM scores, computed using the spp package [11]; 2) the QC tag, which is a discretized version of the RSC score as described in [4]; 3) the recently proposed RSC, NSC based on the Jaccard Index of the strand-shift analysis, and the background uniformity (bu) metrics computed by the SSP package [7]; 4) the “fraction of reads in the top 1% bins”, which is also part of the GM scores as described above and derived from the “fingerprint plot” originally proposed by the CHANCE tool [8]; 5) the Jensen-Shannon Distance (JSD) computed on the fingerprint plot as implemented in deepTools [22]; 6) in addition to the global density QC indicator (denQCi) and the QC-STAMP scores at 5% dispersion as computed by the NGS-QC Generator tool [21]. The latter two are another example of metrics examining the global read distribution across genomic bins. These are metrics incorporated into the QC-STAMP, which has been proposed as a generalizable score to assess the clustering of reads in multiple types of high throughput sequencing based experimental techniques, thus not specifically limited to ChIP-seq [21].

Thus, we compared our ChIC RF-score against a total of 10 alternative quality control metrics obtained from methods or tools described in 6 different publications. This extensive benchmarking confirmed that the ChIC RF-score outperforms all of the previously proposed by resulting in a larger AUC for all of the four classes of profiles (TFs, Sharp, Broad and RNAPol2) (Fig. 3a, Supplementary Figure S3a). For the denQCi and QC-STAMP metrics the same observations are confirmed with 2.5% and 10% dispersion as well (Supplementary Figure S3b). We also examined the performances on selected individual histone marks (H3K9me3, H3K27me3 and H3K36me3). For H3K9me3 and H3K36me3 the ChIC RF score outperforms all of the other QC (Supplementary Figure S3c and S3d). For H3K27me3 instead it shows a performance comparable to other scores as 6 of them have AUCs within the interval 0.68±0.03. Taken together, these results confirm that our machine learning-based combination of quantitative features can better discriminate ChIP-seq samples quality across several types of marks.

**Figure 3:**
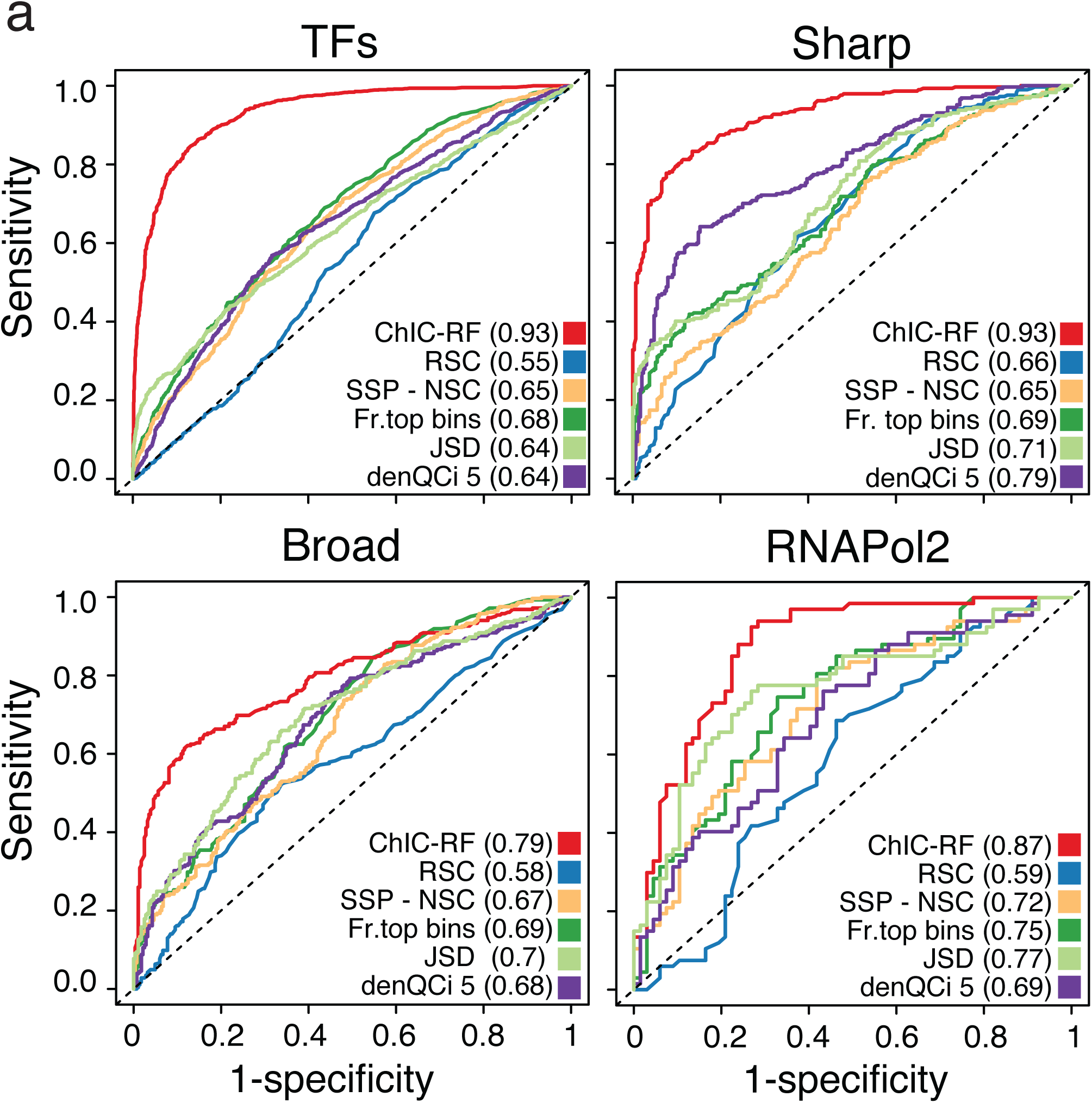
Benchmarking against previous metrics. (**a**) ROC curves summarizing the performance of QC-metrics to assess the quality of ChIP-seq data. The ROC curves are calculated on ENCODE samples versus their simulated problematic counterpart (see methods), grouped into Sharp (287 samples), Broad (285 samples), RNAPol2 (67 samples) and TF (1125 samples) classes. For each class the actual number of samples is doubled by the simulated problematic counterparts. Here we compare the ChIC RF-score; the RSC values derived from cross-correlation strand-shift analysis; the NSC based on the Jaccard Index of the strand-shift analysis computed by the SSP package; the “fraction of reads in the top 1% bins”, which is also part of the GM scores; the Jensen-Shannon Distance (JSD) computed on the fingerprint plot as implemented in deepTools; the global density QC indicator (denQCi) computed at 5% dispersion by the NGS-QC Generator tool. For each score the AUC is indicated in parenthesis in the colour legend.

### Application to test cases

In addition to provide a single score (RF-score) summary, our ChIC framework can also leverage the compendium of QC-metrics as a reference for direct comparison to new samples focusing on individual metrics. To illustrate this feature, we considered a test case with three examples including Sharp, RNAPol2 and Broad marks (Fig. 4). These are samples that have been flagged by ENCODE as potentially problematic and previously excluded from our reference compendium (see methods). For each of these samples, the RSC metric is plotted over the distribution of RSC values across the reference compendium (Fig. 4a), grouped by chromatin marks sub-classes (Sharp, RNAPol2 and Broad) for a more precise estimation of the expected distribution. As shown in (Fig. 4a), these samples have RSC values in the lower tail of the distribution of RSC for their respective chromatin mark type. As an alternative diagnostic plot, the local enrichment profile can be inspected by plotting the metagene profile against the expected profile in the reference compendium for the same chromatin mark. The examples in (Fig. 4b) show how this plot can highlight a generally weak enrichment signal in the three problematic samples. Moreover, the weaker than expected enrichment is especially evident where each of these marks is expected to be enriched: *i.e.* around the TSS for H3K4me3 and RNAPol2, and along the gene body for H3K36me3.

**Figure 4:**
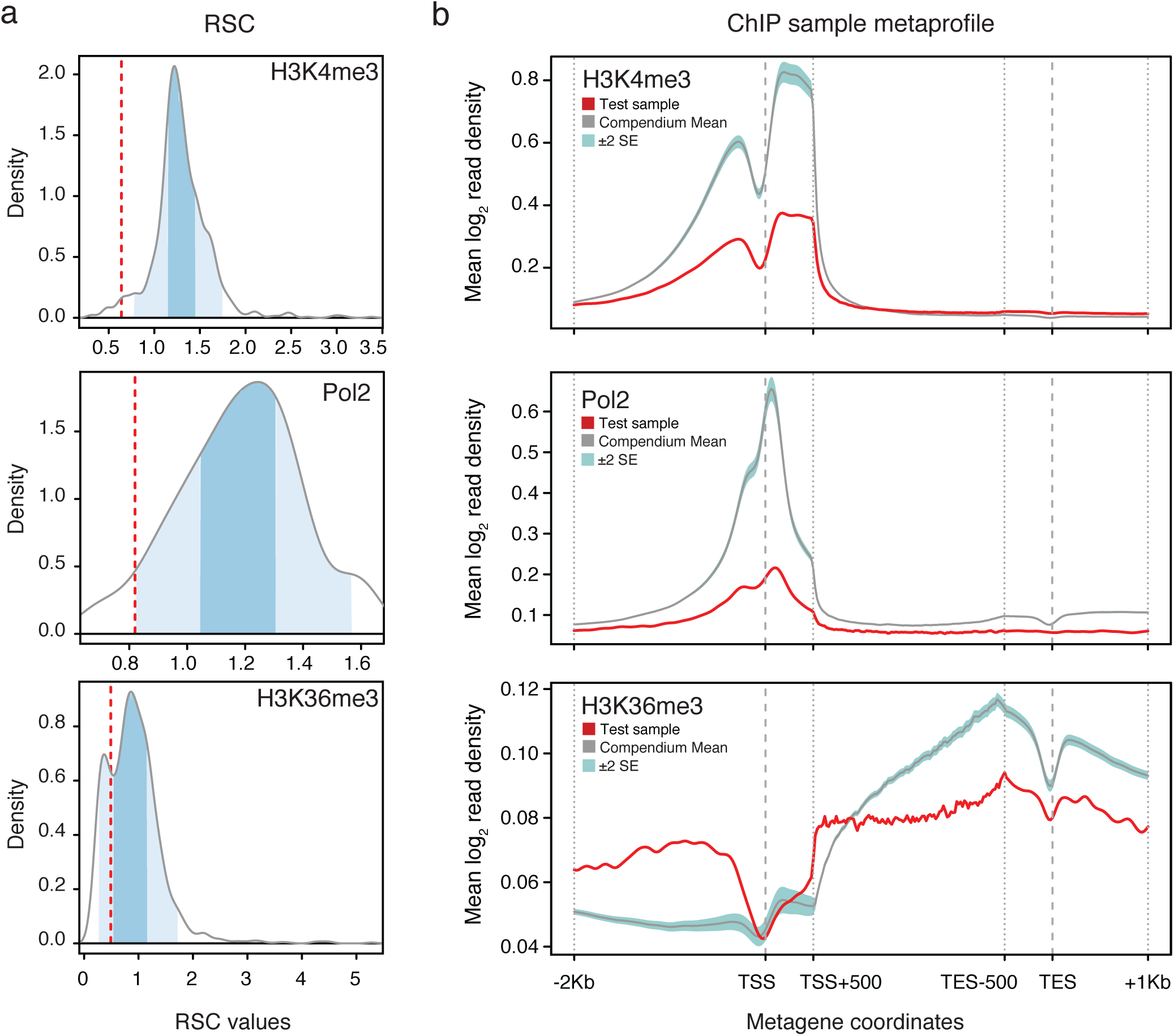
Exploring ChIP-seq quality with the reference compendium. (**a**) QC-metrics of newly analysed ChIP-seq samples can be compared with the reference values of the compendium. Shown is the QC-metric RSC for three problematic ENCODE samples (red dashed line) plotted against the RSC distribution (density plots) of the reference compendium values for the respective chromatin mark. Darker blue shading marks values between 25 and 75 percentiles, lighter blue shading marks values from 5 percentile (lower tail) and up to 95 percentile (upper tail). The problematic examples for chromatin marks H3K4me3 (top), RNAPol2 (centre) and H3K36me3 (bottom) plot2 (IDs: ENCFF000CSO, ENCFF000RPA and ENCFF000BLL, respectively), have been flagged by ENCODE with a red audit as potentially problematic and previously have been excluded by our reference compendium. (**b**) The enrichment pattern for individual samples can be plotted against the expected metagene profile based on data from the reference compendium from published datasets. Here is the average log_2_ transformed normalized (ChIP over input) enrichment signals is shown for three problematic ENCODE samples (red line - same samples as in panel **a**) compared to the compendium’s mean signal (grey line) along with its ±2 standard error interval (blue shadow). The mean signal has been calculated on 369 H3K4me3, 123 RNAPol2 and 287 H3K36me samples.

As a further test case of the ability of our approach to assess sample quality, we analysed a H3K27me3 ChIP-seq dataset in mouse cells with two replicates. While replicate 1 (Rep1) has been successful, the unpublished replicate 2 (Rep2-bad) was affected by problems due to excessive sonication (Supplementary Figure S4a) and a general very low enrichment suggesting a problem in immunoprecipitation as well. After repeating the experiment, a satisfying data quality was obtained also for replicate 2 (Rep2-good). Visual inspection of a known PRC2 targeted genomic region, such as the HoxB cluster, clearly shows that H3K27me3 ChIP-seq signal enrichment is evident for Rep1 and Rep2-good, but not for Rep2-bad (Fig. 5a), for which only an adjustment of y-axis scale would reveal some weak enrichment. The comparison of the three samples to the reference compendium metaprofile also shows that Rep2-bad has a generally low enrichment, thus low quality, compared to the other two samples (Fig. 5b). This confirms the importance of incorporating information on the shape of the enrichment profile, such as the LM features, in evaluating sample quality. Indeed, the difference in samples quality is not equally evident if relying on previously proposed QC strategies. For example, some EM metrics improperly score Rep1 as the worse one (*e.g.* RSC and QC tag) (Supplementary Figure S4b). The ideal QC score should assign a similarly high-quality value to both Rep1 and Rep2-good, while assigning a much lower quality value to Rep2-bad. Indeed, among other alternative metrics included in our benchmarking, our ChIC RF-score stands out for clearly showing a large difference in score between Rep2-bad vs Rep2-good, while at the same time showing a similarly high score for Rep1 vs Rep2-good (Fig. 5c). The RF-score has the largest relative difference between good samples (Rep1 and Rep2-good) vs the bad one (Rep2-bad), if excluding the JSD score that has a large difference between Rep1 vs Rep2-bad, but it also misleadingly assigns a lower score to Rep2-good compared to Rep1, as shown in (Fig. 5c). Instead, by applying our analysis procedure, we immediately and automatically detect a low RF-score for the problematic Rep2-bad sample (0.28), i.e. below the optimal threshold (0.5; Supplementary Table S1). Whereas Rep1 and Rep2-good obtained higher and similar RF-score values (0.86 and 0.85, respectively), confirming their quality is similar and good (Fig. 5a).

**Figure 5:**
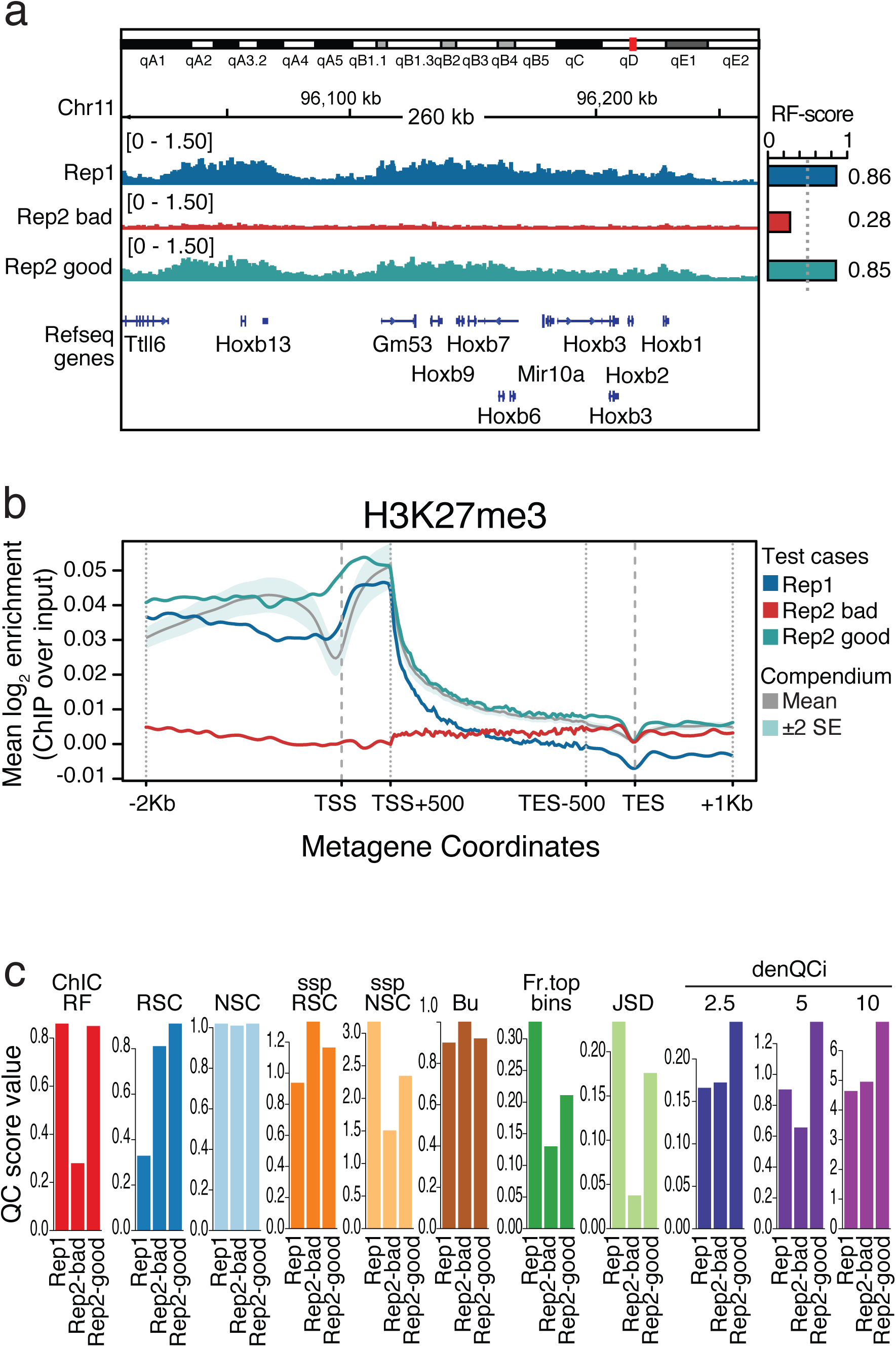
Analysis of a test case. (**a**) ChIP-seq signal for the test case H3K27me3 ChIP-seq samples. A representative region of known PRC2 targets on which H3K27me3 enrichment is expected, i.e. the HoxB genes cluster, is shown. ChIP-seq reads density of good quality Rep1, problematic Rep2-bad and good quality Rep2-good is shown for the region chr11:96000000-96260000 (mm9 genome assembly). The horizontal barplot on the right side shows the RF-score obtained with the H3K27me3 model for each of the samples. The dashed grey line marks the critical threshold value of RF-score discriminating good and poor quality samples. (**b**) The metagene profiles of the test case samples are compared to the reference compendium average metagene profile. The average log_2_ transformed normalized (ChIP over input) enrichment signal is reported. Rep1 (blue) and Rep2-good (green) show enrichment in line with the expected profile, whereas Rep2-bad (red) shows a low enrichment signal. (**c**) Barplots reporting the scores computed by ChIC RF-score and other alternative metrics included in our benchmarking. Each metric has a specific range of values (different y-axis scale), but the relative differences between good samples and failed one can be appreciated and compared across metrics.

We used this as challenging test case as *i)* the problematic sample was affected by a combination of experimental technical issues, as discussed above; *ii)* the organism analysed is mouse and *iii)* H3K27me3 proved to be the most challenging histone mark for all the scoring methods examined in our benchmarking (Supplementary Figure S3c). In this respect, our results confirm the reliability of the single RF-score as a summary of data quality, as well as the applicability of our models to different mammalian datasets. We also tested our framework performance on other non-mammalian datasets, showing that our RF-score can capture subtle differences in ChIP enrichment (Supplementary Figure S5).

### All-in-one: ChIP-seq quality Control package

We implemented all the functionalities and tools presented in this article in a Bioconductor package named “ChIC” (**<u>ChI</u>**P-seq quality **<u>C</u>**ontrol package), with its most updated stable release available at https://bioconductor.org/packages/devel/bioc/html/ChIC.html. The software package is paired with a data package (“chic.data”) containing the reference compendium. Key features of the package making it a unique all-in-one solution for quality assessment include: a single command to compute the comprehensive set of metrics (LM, EM and GM), the machine learning-based single value (RF-score) scoring function, and functions for visualization and summary plots of QC-metrics compared against the reference compendium (Fig. 1a).

It may be worth remarking that ChIC is a package for quality control on ChIP-seq data, thus it’s not designed to call peaks *per se*. However, ChIC relies on a peak-calling algorithm [11] to calculate peak-related QC-metrics (such as FRiP). To compute these and other metrics, ChIC requires a pair of ChIP and input control samples. However, the RF-score calculated by ChIC should be intended as a score for the quality of the ChIP sample, within a pair of ChIP and input samples.

Our ChIC package will be useful for both experts and novices in the field. For non-expert users, the package provides a simple solution to compute a comprehensive set of quality metrics for one or more ChIP-seq samples. The values can then be easily compared to a large reference compendium of data (Fig. 4). For the expert users, the package provides a framework that can be easily incorporated in pre-existing pipelines to assess new ChIP-seq samples, as well as a solution for a quick and automated screening of large amounts of ChIP-seq data from proprietary or public sources alike.

## DISCUSSION

ChIP-seq is a powerful and flexible technique to map TF binding sites as well as chromatin mark distributions genome-wide. Despite being widely adopted, the variability in data quality in public databases [4, 26] and the lack of a generalized consensus on quality assessment parameters are still a cause of concern in the field. Its versatility poses some challenges in assessing objectively whether an experiment was successful or not, because different types of ChIP targets are expected to yield very different enrichment profiles. For these reasons, the expertise of the data curator is important for data quality assessment, thus making it a partly subjective operation.

To address these open problems, we introduce novel quantitative metrics based on the local shape of the ChIP-seq enrichment pattern, as summarized in metagene profiles (LM). While metagene profiles have been previously used to summarize ChIP-seq data, to the best of our knowledge this is the first time that they are used to derive quantitative QC-metrics. The rationale behind this approach is that visual inspection of the ChIP-seq profile is usually an important part of data quality control, yet this is prone to biases, subjective and hardly reproducible. Instead, we use LM to capture and quantify in detail the characteristics of different binding profiles (sharp or broad) for a generalizable and automatic ChIP-seq quality control scoring procedure. It is worth remarking that, compared to all the previous QC methods, ChIC is the only one explicitly considering the expected shape of the enrichment profile of different chromatin marks. This is a crucial point in the rationale of our approach that makes it unique among unbiased automatic ChIP-seq QC scoring solutions. As shown in the results section, several of the previously proposed QC-metrics are biased by the enrichment profile shape and cannot reliably be used to predict data quality. Even recently published solutions computing alternative metrics on the strand-shift profile [7] do not clearly impact the overall performance (Figure 3 and Supplementary Figure S3).

Instead, the ChIC approach is based on a machine learning classifier incorporating both LM and other metrics that are at the basis of previously proposed quality control solutions. As such it’s not surprising that ChIC allows achieving better classification performances compared to other scores as they are *de facto* incorporated and extended in the random forest models. The results obtained with our random forest models show the importance of combining different types of QC metrics by binding characteristics when assessing ChIP-seq quality.

It’s worth remarking that, to quantitatively compare the performances of different quality control methods, the best practice would require the identification of a “gold standard” benchmarking dataset with two large and numerically balanced sets of true positive and true negative quality samples. This is problematic, as the number of failed samples is generally much lower than the successful ones. Moreover, failed samples are usually not publicly released. The challenges in quantitative benchmarking of QC metrics is attested by the fact that, to the best of our knowledge, none of the other tools and procedures for ChIP-seq QC proposed so far in literature provided an extensive quantitative benchmarking, with ROC curves to directly assess the discriminative power for good vs bad quality samples. The results of our benchmarking confirm that the ChiC RF-score is a reliable solution to summarize the sample quality into a single score (Figure 3 and Supplementary Figure S3).

In addition, the comparison of individual samples against the reference compendium, provides easy to interpret information on data quality (Fig. 4). To this concern, user-friendly data visualization functions are also included in our framework. We implemented all these tools in the ChIC Bioconductor package, and an accompanying data package that includes the compendium of reference QC-values. In this study we analysed more than 3000 published ChIP-seq samples by computing a comprehensive set of QC-metrics. Compared to previously published compendia [19, 27], we collected a wider and more diverse set of QC-metrics, including LM, EM and GM. Our compendium contains a broad range of ChIP-seq profiles, including TFs and multiple types of chromatin marks. While other databases previously proposed a compendium of ChIP-seq samples with selected quality scores [19, 20, 27], this is the first time that a reference compendium is directly used to train machine learning models to reliably summarize the sample quality in one single score.

On one hand, ChIC will be useful to non-expert users, who analyse a few samples and may face uncertainties in assessing how much of their own data is in line with expected results. On the other hand, it will also be useful for expert users, who routinely analyse many datasets and may need to dedicate a significant amount of time to screen multiple types of ChIP-seq profiles to remove potential outliers. As such, we believe the proposed framework will prove to be a valuable tool for the community, to facilitate and improve QC of data from a crucial and widely adopted technology in the epigenomics field.

## MATERIALS AND METHODS

### Datasets

Public ChIP-seq datasets (sequencing reads aligned to Human hg19 genome assembly) have been downloaded from ENCODE data portal [28] (https://www.encodeproject.org/files/) in BAM format and from Roadmap data portal [24] in tagAlign format (http://egg2.wustl.edu/roadmap/data/byFileType/alignments/unconsolidated/). After removing ENCODE samples with “red audit flags”, in total we analysed 3370 ChIP and 566 input, for a total of 3936 samples. 1125 TF and 699 chromatin mark datasets were obtained from ENCODE, the latter including histone marks and RNAPol2 samples. 1546 chromatin mark samples were obtained from Roadmap (Table 1). ChIP-seq samples that are not paired with a control sample have not been analysed in this study. The use of a control sample is crucial to adjust for biases in sequencing reads coverage and therefore a pair of ChIP and input aligned read files was a requirement.

The *Drosophila melanogaster* ChIP-seq datasets (Supplementary Figure S6) were obtained from the modENCODE data portal (http://data.modencode.org/) [29].

A set of problematic ChIP-seq data (negative dataset) was created by simulating lower “signal to noise ratio”. It’s worth remarking that we do not apply a “uniform” down-sampling as that would just simulate a lower sequencing depth. While sequencing depth *per se* is important to determine the statistical power of the ChIP-seq peaks calling analyses [30], the most common problems in ChIP-seq experiments are related to antibody specificity, variability in sonication, immunoprecipitation efficiency and various combinations of these issues, that generally result in a lower signal-to-noise ratio in ChIP-seq data. As such, we specifically simulated problematic datasets with lower signal-to-noise ratio while preserving the main peaks localization. This was achieved by first doing a coarse grain call of regions with ChIP over input control enrichment (using “get.broad.enrichment.clusters()” function in the spp package [11]) then down-sampling by 60% the reads mapping within these regions. With this approach, we simulated a problematic counterpart for each reference sample.

### Mouse test case dataset

Mouse muscle satellite cells suspension was fixed in 1% PFA solution for 9 minutes and quenched with 125 mM Glycine before FACS sorting. FACS-isolated satellite cells were resuspended in SDS buffer (100 mM NaCl, 50mM TrisHCl pH 8.1, 5 mM EDTA, 0.2% NaN3, 0.5% SDS) and stored at −80°C (100 μL/106 cells). For ChIP analysis 2-2.5 million fixed cells (collected from 6 to 9 mice) were used for each experiment. Cells were thawed on ice, resuspended in fresh SDS buffer and incubated at 4°C in mild agitation for 3 hours, passing them through a 0.50×16mm syringe needle every hour. The solution was then adjusted to IP Buffer composition (100 mM Tris pH 8.6, 0.3% SDS, 1.7% Triton X-100, 5 mM EDTA) and cells were sonicated with Branson Digital Sonifier to shear chromatin to target average of 200 bp fragments. The 2% of the total volume from each sample was taken as input chromatin. The remaining fragmented chromatin was incubated with 1 mM PMSF (Sigma, 93482) and 4 μg of the H3K27me3 antibody (Millipore 07-449) on a rotating wheel overnight at 4°C. The next day, protein G beads (Life Technology, 1004D) were added (80 μL) and samples were incubated for additional 2 hours on the rotating wheel at 4°C. The beads were washed with Low Salt solution (150 mM NaCl, 20 mM TrisHCl pH 8.0, 2 mM EDTA, 0.1% SDS, 1% Triton X-100), High Salt solution (500 mM NaCl, 20 mM TrisHCl pH 8.0, 2 mM EDTA, 0.1% SDS, 1% Triton X-100), Low Salt solution and then TE NaCl (50 mM NaCl, 10 mM TrisHCl pH 8.0, 1 mM EDTA). Crosslinking was reversed by incubating the beads at 65°C overnight in Elution buffer (50 mM TrisHCl pH 8.0, 10 mM EDTA, 1% SDS). Crosslinking was also reversed. in Input chromatin in Elution buffer overnight at 65°C. The next day, all samples were diluted with one volume of TE 10:1, treated with 0.2 ug/mL RNase A for 2 hours at 37°C and then with 0.2ug/mL Proteinase K (Sigma P2308) for 2 hours at 55°C. DNA was isolated through standard phenol/chloroform extraction, followed by precipitation and resuspension in 31 μL of 10 mM Tris-HCl pH 8.0. Libraries were created using Biomek FX automatic liquid handler (Beckman Coulter), then qualitatively and quantitatively checked using Agilent High Sensitivity DNA Kit (Agilent Technologies, 5067-4627) on a Bioanalyzer 2100 (Agilent Technologies). Libraries with distinct adapter indexes were multiplexed and, after cluster generation on FlowCell, were sequenced for 50 bases in the single read mode on a HiSeq 2000 sequencer. Sequencing reads were aligned to the mm9 mouse reference genome using bwa aln (version 0.7.12) with options -n 2 -k 2 and saved in sam format with bwa samse. The sam files were converted to bam and coordinate sorted with samtools (version 1.3.1).

### Chromatin mark classes

ChIP-seq profiles are grouped into TFs and chromatin marks. When indicated in the text, we further sub-group chromatin mark samples by the expected shape of their enrichment peaks into “Sharp” and “Broad” [3, 31] (Table 2) following the ENCODE3 guidelines (https://www.encodeproject.org/chip-seq/histone/). We also define an additional sub-class for RNAPol2 samples, as it is expected to show a peculiar mixed peak shape profile. Indeed, in our analyses we observe marked differences in the metrics computed for RNAPol2 samples.

### Metagene profiles

Metaprofiles show the average ChIP enrichment over multiple genomic regions of interest. When they are centred over genes, they are also referred to as “metagene profiles” [16]. To compute metagene profiles we considered RefSeq transcripts (hg19 genome version) downloaded from UCSC Genome browser (Jan 4, 2016 build) [32]. For each gene locus, the most upstream and downstream start and end coordinates of its annotated transcripts are used. Genes are filtered if shorter than 2Kb or overlapping other genes on the same or opposite strand. To avoid interferences, we also removed genes overlapping each other in the extended window (2Kb upstream and 1Kb downstream of the gene) considered for metagene profiles.

Metagene profiles are constructed using either the normalized (ChIP over input control) enrichment signal or the read coverage for the ChIP sample considered alone. For ChIP sample considered alone we used the spp package function *get.smoothed.tag.density()* [11] for Gaussian kernel smoothing of reads distribution, to estimate their density genome-wide. The read densities are then normalized by library size and log_2_ transformed (after adding a pseudocount to avoid log transformation of zero coverage values). For “ChIP over input” the Gaussian kernel smoothing is performed for ChIP and input control separately, then normalized by library size and their log_2_ ratio is computed after adding the same pseudocount to both samples.

Two types of metagene profiles are constructed: (1) the unscaled single-point metagene and (2) the scaled whole gene metagene.

In the unscaled single-point metagene, the annotated TSS or annotated transcript end (TES) are used as central point (Supplementary Figure S1c). The direction of transcription is considered so that around the TSS the positive coordinates correspond to transcribed regions and negative coordinates correspond to upstream flanking regions. The opposite applies for TES centred plots. For both profiles we used equally sized bins (201 bins of size 20bp) covering a +/-2Kb window around the mid-point. Considering all genes for each bin we computed the average ChIP-only signal or ChIP over input ratio.

In the scaled whole gene metagene encompassing the entire gene body including 2Kb upstream (promoter) and 1Kb downstream flanking regions. The flanking regions as well as the first and final portions (500bp each) of the gene body are represented with real (unscaled) distances, by dividing these regions in equally sized bins (301 bins of size 20bp).

The central portion of the metagene profile is scaled to fit a common size (Fig. 1a). This is achieved by dividing the gene body into a fixed number of bins (100 bins but variable size). For each bin we computed the average ChIP-only signal or ChIP over input ratio, as defined above.

The metagene profiles used to compute LM scores are based on all the annotated gene loci filtered only by size and distance, as described above. The list of all genes will include both active and silent ones. As such, for histone marks associated to active chromatin the average enrichment signal in the metaprofile will not be as high as if genes were filtered by transcriptional activity, or vice versa for chromatin marks associated to silenced regions. However, consistently using a comprehensive list representing all annotated genes (both active and inactive) allows unbiased detection of the profile shape for active and inactive chromatin marks alike, which is the purpose of the LM.

### Local enrichment profile Metrics (LM)

The use of two different types of metagene profiles (TSS/TES single point and whole gene body profiles) is important as different types of chromatin marks can be enriched at promoters or over the gene body. The combined use of both metagene types allows capturing more details of the local shape of enrichment profiles. For each ChIP experiment we used a total of 6 metagene profiles: i.e. 3 profile types (TSS, TES and whole gene body profiles) each of them in 2 versions (ChIP-only or ChIP over input signal). From these we derived a total of 214 quantitative metrics, so as to start with a comprehensive set of quantitative features (capturing all details of the enrichment profile shape), which then will be filtered before feeding them into the machine learning classifier.

For the TSS metaprofiles the computed metrics include: local maxima (x, y coordinates), cumulative sum (area) of the enrichment profile values, metaprofile variance and quartiles (i.e. 0, 25, 50 and 75 percentiles), all of them computed considering multiple regions within the +/-2Kb window centred on TSS (Supplementary Figure S1c). In total we collected 43 quantitative metrics for the metaprofiles around the TSS and TES, each of them considering both the ChIP-only or ChIP over input normalized profiles, in total 172 features per sample. For the whole gene body metagene we collect additional 42 QC-metrics per sample from the ChIP-only and ChIP over input profiles (Fig. 1a). See (Supplementary file 2) for the complete list of LM.

### ENCODE Metrics (EM)

For computing EM values we followed the procedures described in [3] with a few modifications. Namely, we retrieve the following values concerning library size and complexity: the total number of uniquely mapped reads, uniquely mapped reads corrected by the library size, mapped reads (including non-uniquely mapped ones); the read length; the fraction of non-redundant mapped reads (NRF), i.e. the ratio between the number of uniquely mapped reads after removing duplicates divided by the total number of mapped reads (unique+non-unique); the NRF adjusted by library size or ignoring the strand direction; the PCR-bottleneck coefficient PBC (number of genomic locations where exactly one uniquely mapping read is found, divided by number of genomic locations to which at least one read maps uniquely). We also consider the number of called binding positions (based on the enrichment of the ChIP over control sample) using the WTD method in the spp package [11](*find.binding.positions()* function with FDR of 0.01 and with e-value of 10). We compute metrics based on ChIP-seq peaks, including the fraction of usable reads in the peak regions (FRiP) [3], for which we call sharp peaks (*add.broad.peak.regions()*[11]) and broad peaks (get.broad.enrichment.clusters()[11]) to obtain two types: the FRiP_sharpPeak and the FRiP_broadPeak. Both FRiP variants are always calculated, independently of the expected peak shape.

Additional EM values are based on cross-correlation (CC) profiles of reads mapping positions on positive vs negative strands [3], as implemented in the spp package [11, 17] and described in [4]: the coordinates of the main peak in the CC profile (fragment-length strand shift value on x-axis, and CC coefficient value corresponding to the height of point A on y-axis); the coordinates of the “phantom-peak” [3], i.e. the peak with strand-shift (x-axis) corresponding to the sequencing reads length and CC coefficient value equal to height of point B (y-axis); the baseline of CC coefficient values at extreme strand-shifts (height of line C on the y-axis). Using these values we also derive the Normalized Strand CC Coefficient (NSC) defined as the ratio NSC=A/C, and the Relative Strand CC Coefficient (RSC) defined as RSC=(A-C)/(B-C) [3] (Supplementary Figure S1a). The “quality control tag” is obtained by discretizing the RSC values [4]. In total, we consider 22 metrics in this category (Supplementary file 2).

### Global enrichment profile Metrics (GM)

GMs are derived from the analysis of the cumulative distributions of reads coverage per genomic bin in each pair of ChIP and control samples. This plot of cumulative distributions was originally proposed as a diagnostic feature in the CHANCE tool, implemented in MATLAB [8] and in the deepTools package as “plotFingerprint” [22]. We extended this approach by extracting specific quantitative features from the plot. Namely, we partition the genome in equally sized genomic bins and count the read coverage for each bin. Bins are then ranked by coverage and the cumulative distribution is plotted with the fraction of the total coverage on the y-axis and the fraction of total number of bins on the x-axis (Supplementary Figure S1b). From the plot we derive 9 quantitative metrics (Supplementary file 2) including the (a) fraction of bins without reads (zero coverage) for ChIP and input; (b) point of maximum distance between ChIP and input cumulative distributions (including the x,y coordinates for the points on ChIP and input cumulative distributions, the absolute difference between the two y-coordinates, and the sign of the difference); (c) fraction of reads in the top 1% of bins with highest coverage for ChIP and input samples considered separately.

### Benchmarking against other tools

Strand-shift profile (SSP) [7] package is a tool for quality assessment of ChIP-seq data without peak calling and is based on a strand-shift profile computed using the Jaccard-Index on the positions of reads mapped on positive and negative strands. SSP provides QC metrics like the NSC (SSP-NSC), the RSC (SSP-RSC) and the background uniformity (Bu). We run SSP (v1.1.2) on our compendium of ENCODE datasets and simulated negatives (Table 1) and assessed the discriminative ability of its proposed QC-metrics in the ROC curves.

On the same datasets we also run the “plotFingerprint” (v 3.1.3) function available in deepTools package [22] to calculate the Jensen-Shannon distance (JSD) between each ChIP and input pair. The JSD value ranges from 0 to 1 indicating the difference between the two curves and is conceptually related to the GM scores included in our compendium.

Another set of metrics assessing the global distribution of reads include the QC-STAMP related metrics that we computed using the NGS-QC Generator (v0.31) [21]. This tool computes the global density QC indicator (denQCi), which estimates the robustness of the ChIP-seq profile based on the subsampling of reads and the difference between the random subsampling target percentage and the actual observed subsampling at each genomic position. Namely we use the denQCi at 50% reads subsampling, as this is used as a reference metric to evaluate profiles in the original study by Mendoza-Parra *et al.* [21]. In particular we considered denQCi for dispersion intervals 2.5%, 5% and 10%, as computed by NGS-QC Generator. As defined in the original article, the NGS-QC Generator tool also computes the QC-STAMP, defined as the ratio between the denQCi score and the similarity QC score from the same algorithm. Thus, we also considered the QC-STAMP score at dispersion intervals 2.5%, 5% and 10% in our benchmarking. It’s worth remarking that in the on-line database by the same authors, the QC-STAMP scores for the three dispersion intervals are computed over a large compendium of published datasets. However, these include not only ChIP-seq, but samples from any high-throughput sequencing-based technique yielding reads enriched in specific genomic locations, such as RNA-seq, CAGE-seq, 4C-seq, FAIRE-seq and several others. In fact, the QC-STAMP method was initially proposed as a generic QC procedure for any high throughput sequencing protocol resulting in reads enriched in specific genomic regions. Thus, not tailored on ChIP-seq specific characteristics. Moreover, in the on-line database, the QC-STAMP distributions are discretized into quartiles and translated into a letter code: with letters from D, C, B and A assigned from the lowest to the highest quartile, respectively. However, the discretization into the letter code is performed by pooling the QC-STAMP scores for every sample in the database, i.e. including non-ChIP-seq samples, which may have quite different distributions. To be able to construct a ROC curve and to calculate an AUC for the benchmarking we used the QC-STAMP scores in their continuous non-discretized version (i.e. the numeric values, not the letters encoding).

### Feature correlation and principal component analysis

Each ChIP-seq sample is described by a comprehensive set of 245 quantitative features (QC-metrics), including LM, EM and GM. To remove uninformative features before training the machine learning models, we applied a feature selection procedure similar to [33] on each set of QC-metrics separately. After standardizing each QC quantitative feature (subtracting mean and dividing by standard deviation) for EM, GM and LM separately, we applied hierarchical clustering (“complete linkage”) based on the distance measure *d* = 1 − |*r*| where *r* is the Pearson correlation coefficient between pairs of features across the reference values in the matrices. Trees are cut at 0.1 and for each cluster we chose the medoid as representative feature (Fig. 3a, Supplementary Figure S2a). A medoid is defined as the most central data point in the cluster, where its average distance from all the data points in the cluster is minimized. The procedure has been applied separately on TFs ChIP-seq samples (1125) and chromatin marks samples (2245), finally reducing the features to 80 for the chromatin marks and to 86 for the TFs datasets.

After scaling and centring the values, the *prcomp()* function in R was used to perform principal component analysis (PCA) with the reduced set of features (Supplementary Figure S2b).

### Random forest training and prediction

Assessing data quality can be described as a binary classification problem with the QC-metrics as independent input variables and the data quality as the dependent output variable (class labels: good or poor quality). We used random forest method (“caret” R package [34]) and tuned the parameter “*mtry*” (number of features randomly sampled as candidates at each split) [35] in a 10-fold cross-validation using ENCODE data as positive (+1) and the simulated failed counterparts, generated as described above, as negative (−1) class. The “*mtry”* value with the highest F1-score was used for the final training of the model. The F1 score is a balanced performance measure used to assess the predictive ability of a classifier. The score ranges from 0 to 1 (0 for the worst prediction and 1 for perfect one). The Roadmap samples and their simulated failed counterparts were used as an independent set to test the performance of the classifiers calculating the ROC curve (with sensitivity and specificity) and the corresponding area under the curve (AUC) (Supplementary Table S1). We created separate models for TF and chromatin mark datasets where chromatin marks were further divided into “RNAPol2”, “Sharp” and “Broad” subclasses, as described above (Fig. 3b). Additional models were made for individual chromatin marks (H3K9me2, H3K27me3 and H3K36me3), as described in the results section (Fig. 3c). For each model, we determined the individual feature importance scores provided by the “caret” package during random forest model generation.

## DECLARATIONS

### Ethics approval and consent to participate

Not applicable

### Consent for publication

Not applicable

### Data availability

ChIC R/Bioconductor package is open source with the latest version available at the URL: “https://bioconductor.org/packages/devel/bioc/html/ChIC.html“. The accompanying ChIC.data package, containing the reference compendium data and additional sample data is available at the URL “https://bioconductor.org/packages/devel/data/experiment/html/ChIC.data.html”. Additional file 1 and 2 contain supplementary figures and tables. Raw data of the mouse test case is available in GEO database with accession number GSE119932 (for Rep2-bad sample) and GSE123724 (for Rep1 and Rep2-good samples).

### Author’s contributions

CML implemented the method and performed the analysis with the aid of FF. IT, KP, ES and FF provided support in the development of the Bioconductor packages. IT collaborated in running the analyses and fixing code. FL, AB, SV and CL provided the ChIP-seq test case experiment. FF designed and supervised the work. CML and FF wrote the manuscript. All authors provided critical feedback and helped shape the research, analysis and manuscript. All authors read and approved the final manuscript.

## Acknowledgements

We thank Laura Riva (Sanger Institute), Marco Morelli (San Raffaele Hospital), Oriana Romano and Silvio Bicciato (UNIMORE) for critical feedback on earlier versions of the manuscript. We also thank Peter Kharchenko (HMS) for specific spp package related functions (as detailed in the ChIC package source code) and Eugenia Galeota (IIT) for help with C code issues. We thank CINECA for computing resources provided in the context of ISCRA-C project “CONCEPt”, and in particular Giovanni Chillemi and Tiziana Castrignanò for their advice and support.

## SUPPORTING INFORMATION LEGENDS

**Supplementary File 1** (Livi_etal_SupplementaryFile1.pdf). Document containing Supplementary Figures and Supplementary Table S1 along with their legends.

**Supplementary File 2** (Livi_etal_SupplementaryFile2.xlsx). Complete list of quantitative features computed by the ChIC framework.

## Supplementary Figures

**Figure S1:**
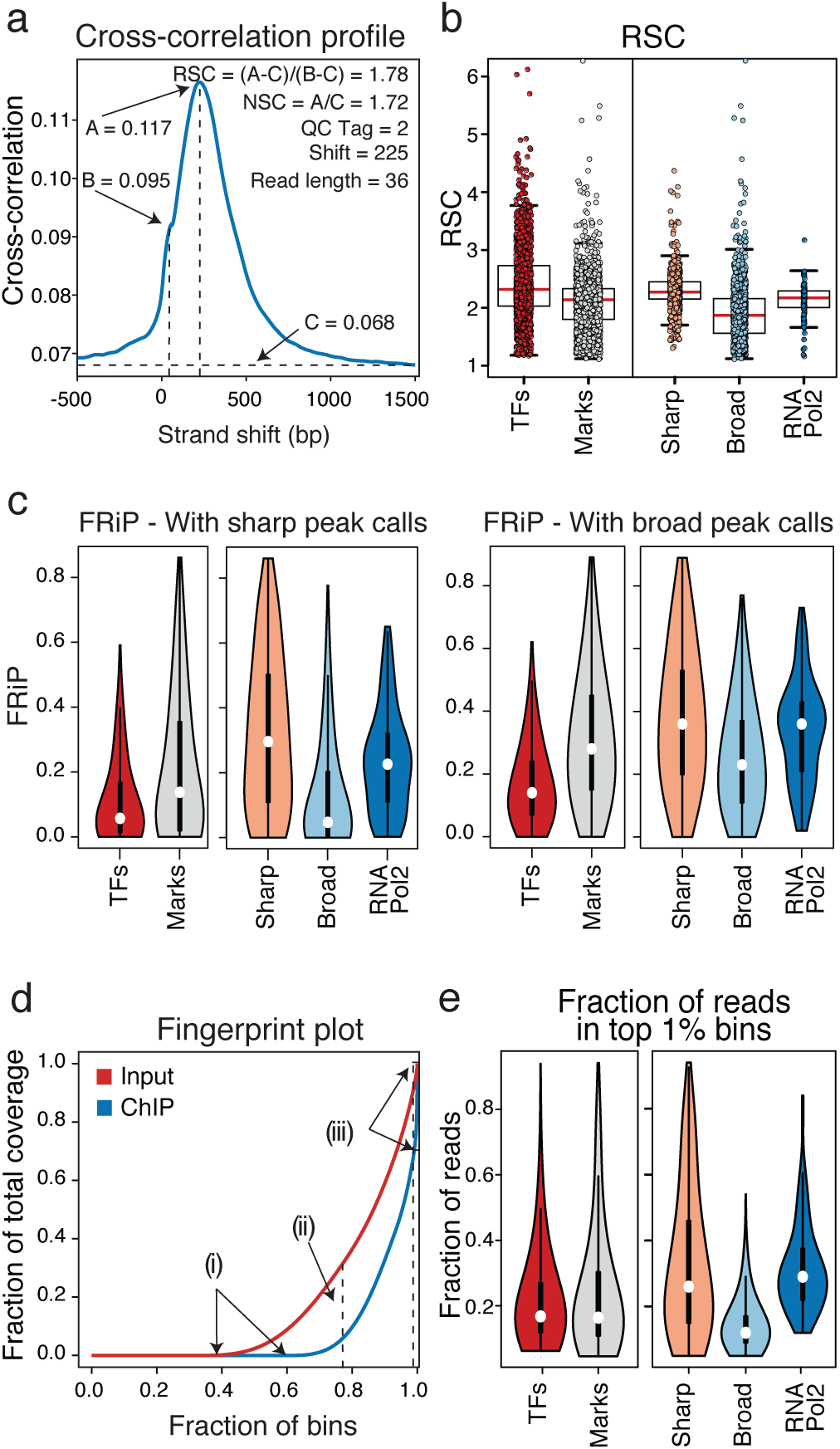
EM and GM scores to assess ChIP-seq data quality. **(a)** Example of strand cross-correlation. Different QC-metrics can be derived from the cross-correlation profile: the normalized strand coefficient NSC, the relative strand coefficient RSC, the strand shift with the highest cross-correlation value (A value and Shift parameter) and the estimated read length (strand shift of point B), as indicated in the plot showing a representative CTCF sample (ENCFF000AH). (**b**) The box plots report the EM “Relative Strand Correlation Coefficient (RSC)” value for different groups of ENCODE and Roadmap data as indicated. Values for TFs (TFs, 1125) and all chromatin marks (Marks, 2245) are shown in the left plot. On the right, RSC values for chromatin marks are subdivided into 3 classes associated to Sharp (941), Broad (1177) and RNAPol2 (127, containing different subunits) profiles. Individual data points are shown over the boxes. (**c**) The violin plots show the distribution of the EM “Fraction of reads under sharp peaks” (left) and “Fraction of reads under broad peaks” (right) for TFs (1125 samples) and chromatin marks (2245) from ENCODE and Roadmap data. The left plots show the distribution of values for the chromatin marks group (see methods), when split into Sharp (941 samples), Broad (1177 samples) and RNAPol2 classes (127), respectively. (**d**) Example of fingerprint plot, i.e. the cumulative distribution of the read counts across genomic bins for input (red line) and ChIP (blue line). The global enrichment profile of a sample can be quantified by taking GM like (i) the fraction of bins without reads, (ii) the point of maximum separation between ChIP and input cumulative distributions, and (iii) the fraction of reads in the top 1% bins (here shown for the ChIP). (**e**) Violin plots showing the distribution of the GM “Fraction of reads in top 1% bins” of ENCODE and Roadmap data for TFs (TFs) and chromatin marks (Marks) in the left plot. On the right the plot shows the value distribution of chromatin marks divided by profile classes (Sharp 941, Broad 1177 and RNAPol2 127, containing different subunits).

**Figure S2:**
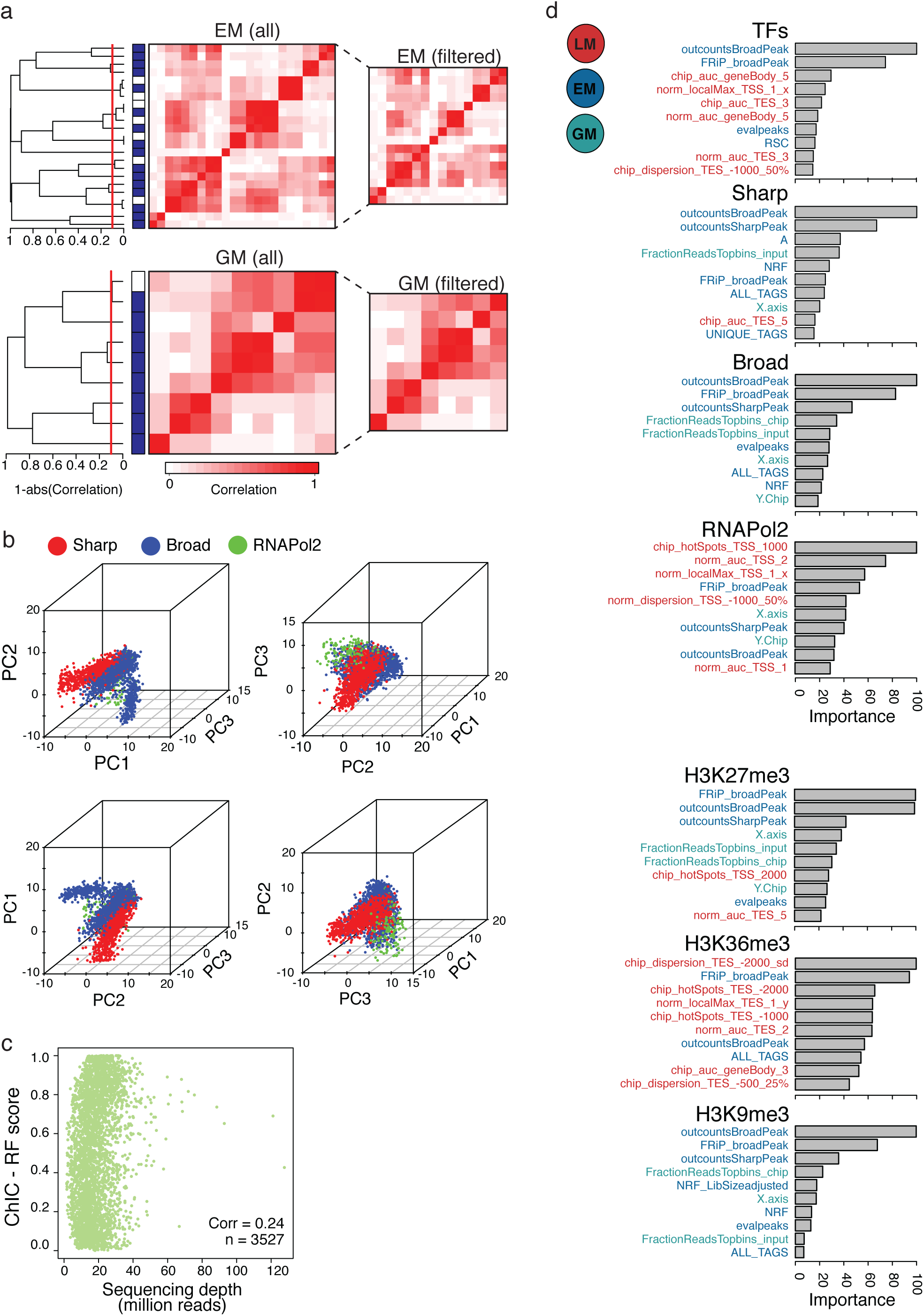
Features selection and importance. **(a)** Correlated features are filtered using hierarchical clustering algorithm on the QC-metrics grouped by category: EM (upper heatmaps), GM (bottom heatmaps) and LM (main text - Figure 2a). The clustering is performed on chromatin marks and TFs (not shown) separately. The dendrogram shows the result of the complete linkage clustering for chromatin marks with distance measured as 1 - (absolute value Pearson correlation). The dendrogram is cut at 0.1 and for each resulting cluster the medoid (blue mark in the sidebar) is retained in the set of filtered features. The left heatmaps show the pairwise Pearson-correlation between all features (before filtering) whereas the right side heatmaps summarizes the correlation of the reduced set of features after filtering. **(b)** Three dimensional dot plots of the first three principal components (PC) of the principal component analysis performed on the reduced set of features. The plots show chromatin mark data from different perspectives and coloured by peak profile classes Sharp (red), Broad (blue) and RNAPol2 (green). (**c**) Scatter plot showing the relation between our RF prediction score (y-axis) and the sequencing depth (x-axis) for all positive and negative samples. The low correlation of 0.24 and the visual examination indicate that there is no bias of the predictor towards the sequencing depth (**d**) Top 10 features ranked by importance for each random forest model. The bar plots show the importance score values, with feature names coloured by category: LM, EM and GM (colour legend).

**Figure S3:**
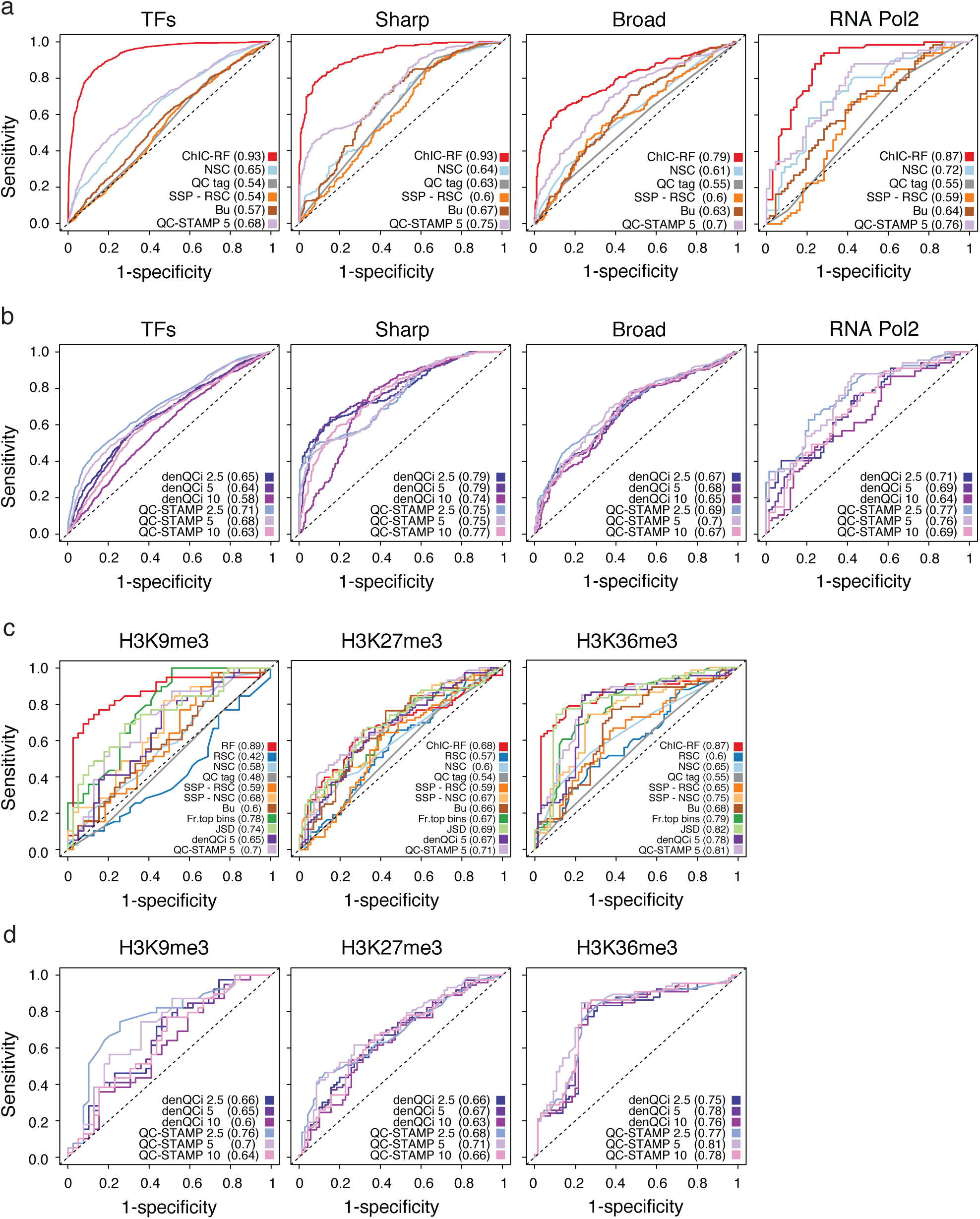
Additional benchmarking results. **(a)** ROC curves summarizing the performance of QC-metrics to assess the quality of ChIP-seq data. The ROC curves are calculated on ENCODE samples versus their simulated problematic counterpart (see methods), grouped into Sharp (287 samples), Broad (285 samples), RNAPol2 (67 samples) and TF (1125 samples) classes. For each class the actual number of samples is doubled by the simulated problematic counterparts. Here we compare 1) ChIC RF-score; 2)the RSC values derived from cross-correlation strand-shift analysis; 3) the QC tag; 4) the NSC based on the Jaccad Index of the strand-shift analysis and the 5) background uniformity (bu) metric, both computed by the SSP package; 6) QC-STAMP score at 5% dispersion as computed by the NGS-QC Generator tool. For each score the AUC is indicated in parenthesis in the colour legend. (**b**) ROC curves as in panel (a), but comparing the performances of additional metrics by the NGS-QC Generator tool. Namely, for completeness of the benchmarking the global density QC indicator (denQCi) and the QC-STAMP scores were computed at 2.5%, 5% and 10% dispersion rates, as indicated in the colour legend of the ROC curves. All of these variations yield comparable results in terms of AUC. (**c**) ROC curves summarizing the performance of different QC-metrics in discriminating ChIP-seq quality for individual subtypes of Broad chromatin marks: H3K9me3 (39 samples), H3K27me3 (73 samples) and H3K36me3 (66 samples), for each class the actual number of samples is doubled by the simulated problematic counterparts. Here we compare all of the 11 main metrics considered in our benchmarking. Namely: 1) ChIC RF-score; 2) the RSC and 3) the NSC values derived from cross-correlation strand-shift analysis; 4) the QC tag; 5) the RSC and 6) the NSC based on the Jaccad Index of the strand-shift analysis, as well as the 7) background uniformity (bu) metric, all three computed by the SSP package; 8) the “fraction of reads in the top 1% bins”, which is also part of the GM scores; 9) the Jensen-Shannon Distance (JSD) computed on the fingerprint plot as implemented in deepTools; 10) global density QC indicator (denQCi) and the 11) QC-STAMP score, both computed at 5% dispersion by the NGS-QC Generator tool. (**d**) ROC curves as in panel (c), but comparing the performances of additional metrics by the NGS-QC Generator tool. Namely, for completeness of the benchmarking the global density QC indicator (denQCi) and the QC-STAMP scores were computed at 2.5%, 5% and 10% dispersion rates, as indicated in the colour legend of the ROC curves. All of these variations yield comparable results in terms of AUC.

**Figure S4:**
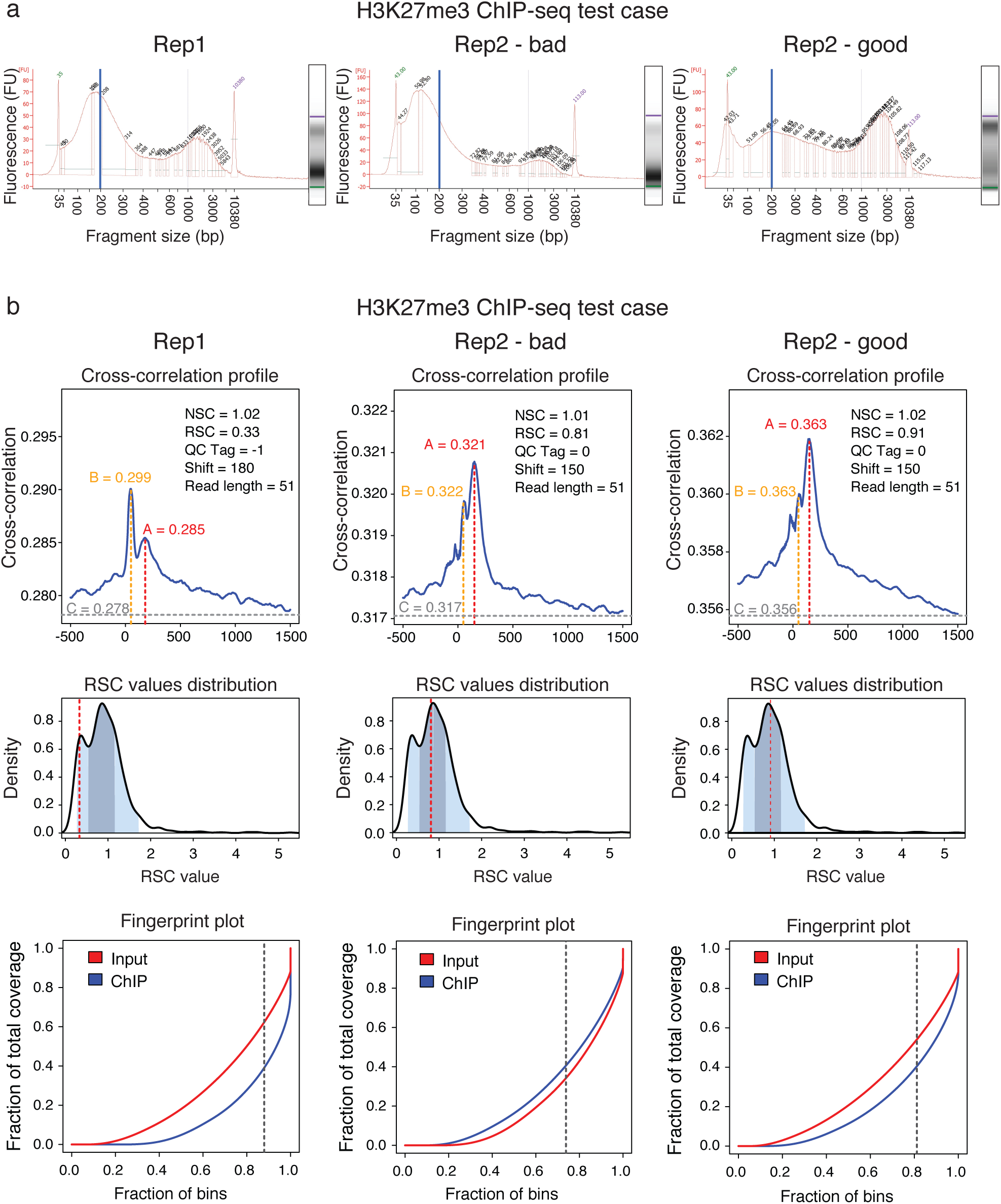
Test case dataset samples. (**a**) Bioanalyzer analysis to assess the quality of the starting material for Rep1, Rep2_bad and Rep2_good. Plots show the DNA size distribution after fragmentation. X-axis shows the fragment size in base pairs (bp) and y-axis the florescence intensity proportional to DNA abundance (FU = florescence unit). The target fragment length was about 200bp (vertical blue line), but “Rep2_bad” fragment size distribution is shifted to the left towards smaller sizes. (**b**) QC diagnostic plots based on EM and GM scores for the test case samples Rep1 (left column), Rep2_bad (centre column) and Rep2_good (right column). Cross correlation profiles (top rows), RSC score (red dashed line) plot against the reference distribution of RSC values (middle row) and fingerprint plots (bottom row) for the three test case samples. Overall, they do not consistently single out Rep2_bad as the real problematic sample.

**Figure S5:**
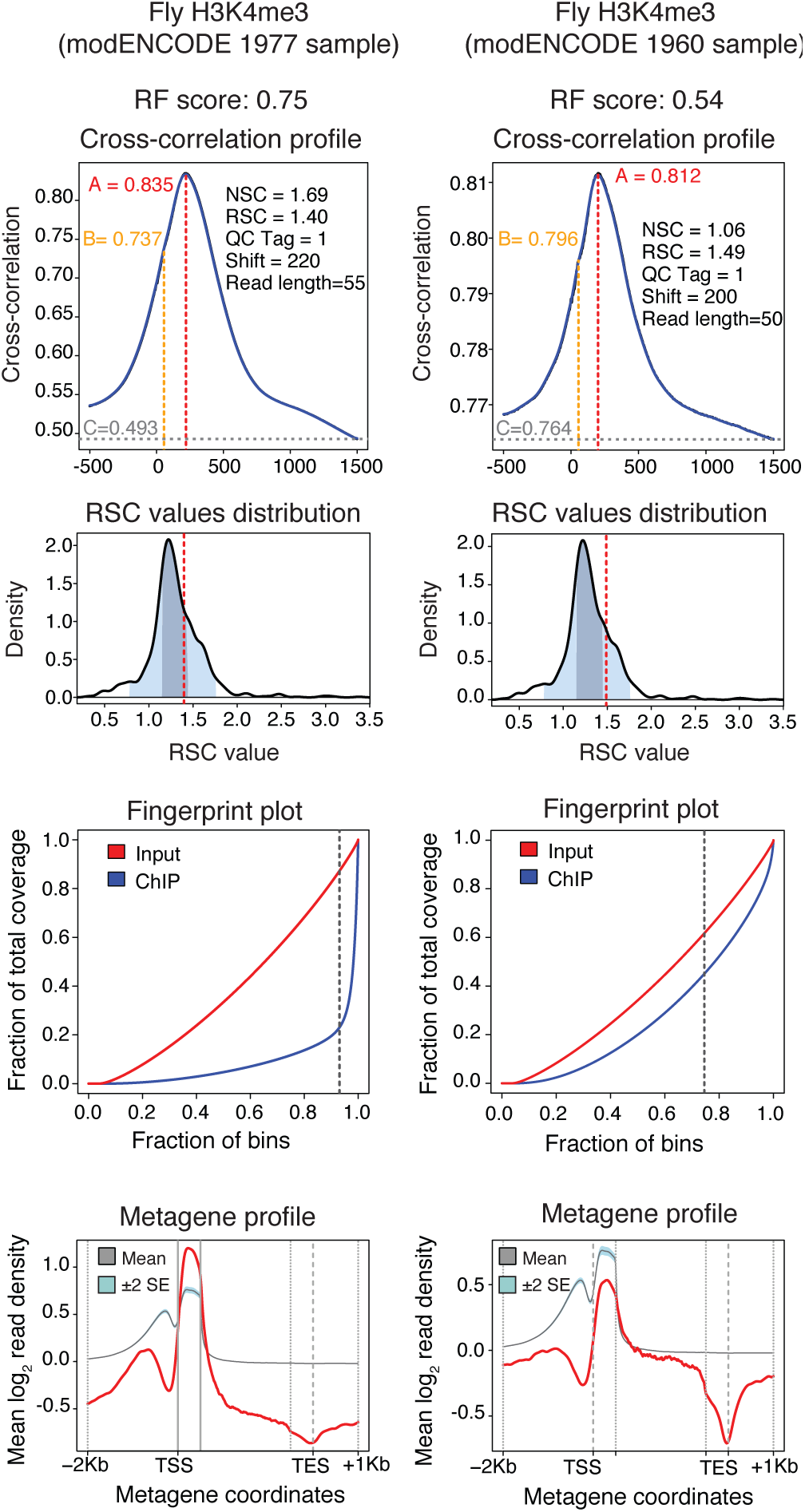
Performance of ChIC on a *Drosophila melanogaster* dataset. The figure reports an example of ChIC performance on other model organisms. Here we show two H3K4me3 samples in *Drosophila melanogaster* 14-16hr embryos from modENCODE: sample IDs Solexa_EL1_H3K4me3.1977 and Solexa_EL7_H3K4me3.1960, in the left and right columns, respectively. The cross-correlation profiles (top rows) and the RSC score (red dashed line) plot against the reference distribution of RSC values (second row) do not highlight any variability in the quality of enrichment in the two samples. Instead the fingerprint plots (third row) and especially the enrichment signal in the metagene profiles (last row) suggest some difference with the second sample having lower enrichment (right column). This variability is clearly reflected and summarized also in our RF score, which is 0.75 for the sample with higher enrichment (left column) and 0.54 for the other sample (right column).

## Supplementary Table

**Table S1:**
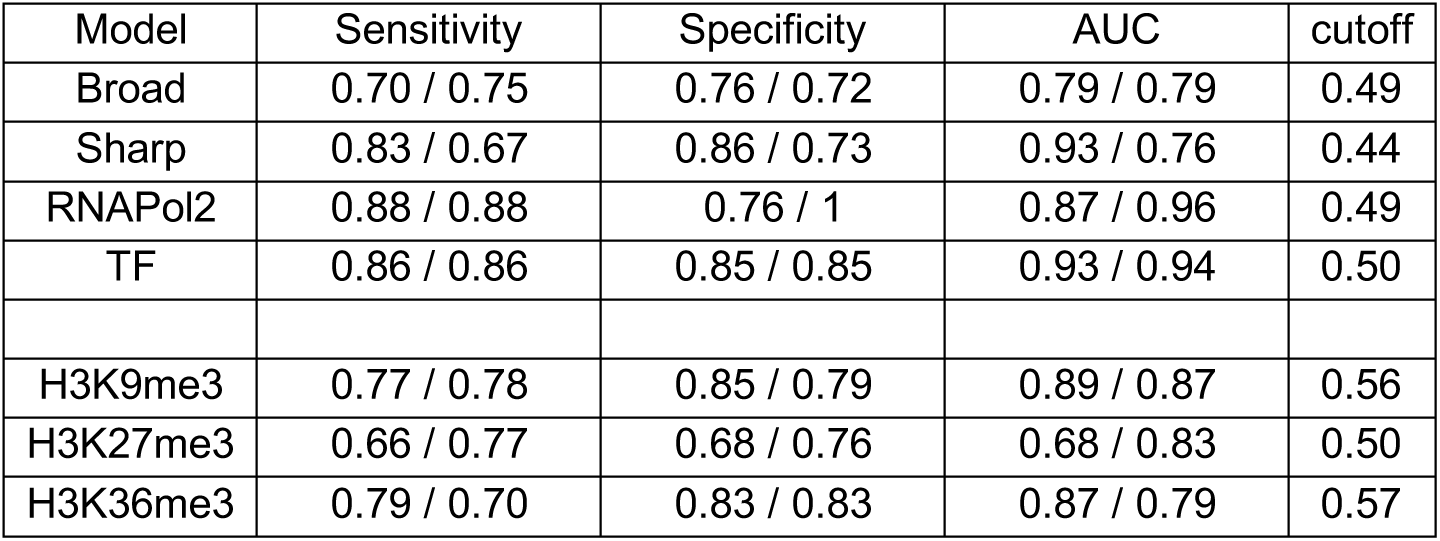
Model performances. The predictive ability of each random forest model is measured by calculating the sensitivity, specificity and the area under the curve (AUC). The sensitivity is given as TP/(TP+FN), the specificity as TN/(TN+FP), with TN being true negative predicted elements, TP true positives, FN and FP being false negative and false positive predicted elements. The AUC is calculated as the area under the receiver operating characteristic (ROC) curve. The optimal cutoff to discriminate positive and negative samples is taken for each model from the respective ROC by minimizing the distance *d* defined as the quadratic sum of *x* and *y: d= x*^*2*^*+(y-1)*^*2*^.

